# Mechanical stretch regulates macropinocytosis in *Hydra vulgaris*

**DOI:** 10.1101/2021.12.03.471193

**Authors:** Taylor D. Skokan, Bert Hobmayer, Kara L. McKinley, Ronald D. Vale

## Abstract

Cells rely on a diverse array of engulfment processes to sense, exploit, and adapt to their environments. Macropinocytosis is a versatile example of such a process, allowing for the indiscriminate and rapid uptake of large volumes of fluid and membrane. Much of the molecular machinery essential for macropinocytosis has been well established. However, most of these studies relied on tissue culture models, leaving the regulation of this process within the context of organs and organisms unresolved. Here, we report that large-scale macropinocytosis occurs in the outer epithelial layer of the cnidarian *Hydra vulgaris*. Exploiting *Hydra*’s relatively simple body plan, we developed approaches to visualize macropinocytosis over extended periods of time in living tissue, revealing constitutive engulfment across the entire body axis. Using pharmacological perturbations, we establish a role for stretch-activated channels, including Piezo, and downstream calcium influx in inhibiting this process. Finally, we show that the direct application of planar stretch leads to calcium influx and a corresponding inhibition of macropinocytosis. Together, our approaches provide a platform for the mechanistic dissection of constitutive macropinocytosis in physiological contexts and reveal a role for macropinocytosis in responding to membrane tension.

## Introduction

Engulfment processes are essential for cells to sense and interact with their extracellular environments. In contrast to receptor-mediated endocytosis and phagocytosis, which rely on interactions with specific ligands, macropinocytosis provides a mechanism to indiscriminately engulf large volumes of extracellular fluid and plasma membrane in the absence of a defined target [1]. This unbiased strategy renders macropinocytosis a highly versatile process, illustrated by the diverse functions it serves (reviewed in [2]), including bulk nutrient acquisition [3,4], immune surveillance [5,6], receptor sequestration [7], and membrane retrieval [8,9]. Regardless of the specific cellular functions served by macropinocytosis, the process involves local remodeling of the actin cytoskeleton to form membrane protrusions, or ruffles, that subsequently fuse to encapsulate extracellular contents. During this process, formation of membrane ruffles depends on the temporally and spatially restricted interactions of phosphoinositides, small GTPases, and actin regulators (reviewed in [10]).

Despite these insights, the factors that initiate macropinocytosis in diverse cell types and physiological contexts are incompletely understood. In some cell types, most notably antigen presenting cells, macropinocytosis occurs constitutively, whereas many other cell types require exogenous growth factor stimulation or pathogen exposure to stimulate macropinocytosis [11,12]. Recent work has identified factors that can enhance the rate of growth factor-mediated macropinocytosis, including excessive cortical localization of the cytoskeleton-membrane linker ezrin [13]. Moreover, recent work in mammalian myotubes revealed that transient reductions in membrane tension following osmotic stress and mechanical stretch can promote growth factor-stimulated macropinocytosis [14,15].

Here, we report the serendipitous discovery of tissue-wide macropinocytosis in the freshwater polyp *Hydra vulgaris*, a representative of the ancestral animal phylum Cnidaria. In contrast to canonical, mammalian models of epithelial macropinocytosis, which require growth factor stimulation (reviewed in [16]), this process unfolds constitutively in *Hydra*’s superficial ectodermal epithelium, accounting for considerable membrane remodeling and fluid uptake. Combining live microscopy with pharmacological perturbations, we establish a role for stretch-activated channels and Ca^2+^ signaling in regulating this process. Finally, by inflating regenerating *Hydra* tissues, we demonstrate a direct role for cellular stretch in regulating macropinocytosis, with high-tension states inhibiting fluid uptake. Together, our findings reveal a role for tissue mechanics in regulating constitutive macropinocytosis and highlight the potential for *Hydra* as a physiological model of macropinocytosis.

## Results

### *Hydra* ectoderm exhibits ubiquitous macropinocytosis

Previous investigations of tissue patterning in *Hydra* have shown that the actin reporter LifeAct-GFP localizes to actin-enriched basal myonemes and apical cell junctions of the outer (ectodermal) epithelium [17,18]. In our studies, we also observed that LifeAct-GFP labeled dynamic, ring-shaped structures that localized to the apical cell membrane (Fig. 1A). Phalloidin staining in fixed, intact animals confirmed that these structures were enriched for actin (Fig. 1B). These actin-rich rings were found at a low frequency along the entire body plan (Fig. S1) and bore a striking resemblance to structures identified in classic scanning electron microscopy of the *Hydra* ectoderm [19]. Since *Hydra* display considerable contractile movement in three-dimensions, we developed a preparation that would allow us to better monitor the dynamics of the actin rings over extended periods. This preparation involved amputating the head and foot from animals and threading the resulting body columns onto filaments to roughly constrain tissue movement to a single plane (Fig. 1C). This approach allowed us to capture the entire lifecycle of actin rings, which formed as an expanding ring, reaching a diameter of 10.49 ± 2.6 μm (n = 35 from 3 independent preparations), and then constricted to a focus and ultimately disappeared (lifespan: 109 ± 32 s; n = 35 from 3 independent preparations; Fig. 1D; Video S1A).

**Figure 1.**
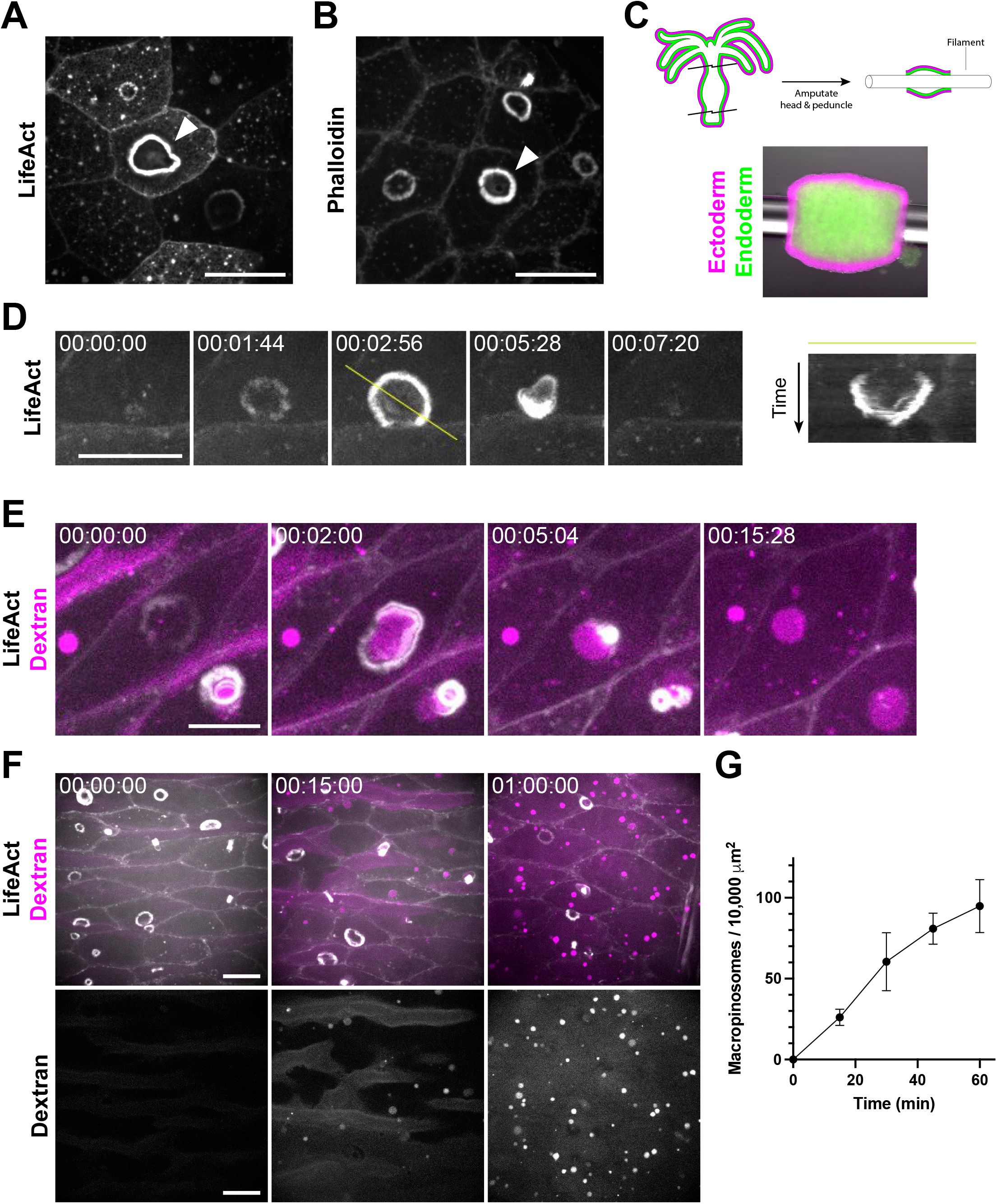
*Hydra* ectoderm exhibits ubiquitous macropinocytosis. A) Representative image of macropinocytic cups (arrowhead) in the ectoderm of a live, LifeAct-GFP-expressing *Hydra*. B) Representative image of macropinocytic cups (arrowhead) in the ectoderm of a fixed *Hydra* stained with phalloidin. C) Schematic (top) and representative image (bottom) of the body column isolation preparation used for prolonged live imaging. Ectodermal and endodermal cells express DsRed2 (magenta) and GFP (green), respectively. D) Representative time-course of macropinocytic cup formation, closure, and dissipation, visualized by LifeAct-GFP (left). Image registration was performed to compensate for translational movement in the body column. Yellow line denotes the axis of the corresponding kymograph showing the complete lifecycle (right). E) Representative time-course of fluorescently labeled dextran (magenta) engulfment by macropinocytic cups, visualized by LifeAct-GFP (white). Image registration was performed to compensate for translational movement in the body column. F) Representative time-course of dextran-filled macropinosome (magenta) accumulation in ectodermal tissues expressing LifeAct-GFP (white). Dextran channel is isolated at (bottom). G) Quantification of dextran-filled macropinosome accumulation (mean ± sd; n = 3 independent sample preparations). (A–F) All frames depict maximum intensity projections of 10–35 μm z-stacks. Time stamps, hh:mm:ss. Scale bars, 20 μm. (D–F) All time-courses depict isolated, threaded body columns.

Given their resemblance to circular dorsal ruffles [20] and macropinocytic cups in mammalian cells ([20]; reviewed in [21]), we considered whether the ring structures may play a role in fluid uptake. To test this, we incubated the threaded body columns in media containing fluorescently labelled dextran. As rings formed, dextran accumulated in the resulting invaginations and was engulfed upon ring constriction (Fig. 1E; Video S1B). The resulting dextran-filled vacuoles persisted within ectodermal cells and accumulated in the tissue over time (Fig. 1F, 1G). Thus, the actin rings denote macropinocytic cups associated with fluid uptake.

### Stretch-activated channel activity regulates macropinocytosis

Intriguingly, we observed a trend of more macropinocytic cups in isolated *Hydra* body columns (0.186 ± 0.140 cups per cell; n = 4 independent preparations) when compared to fixed, intact animals (0.015 ± 0.011 cups per cell; n = 15 from 3 independent preparations). Moreover, the frequency of macropinocytic cups increased over time in body columns following amputation (Fig. 2A; Video S2A). As the incisions required for head and foot amputation are likely to affect tension in the epithelium, we considered whether tissue mechanics may play a role in regulating macropinocytosis.

**Figure 2.**
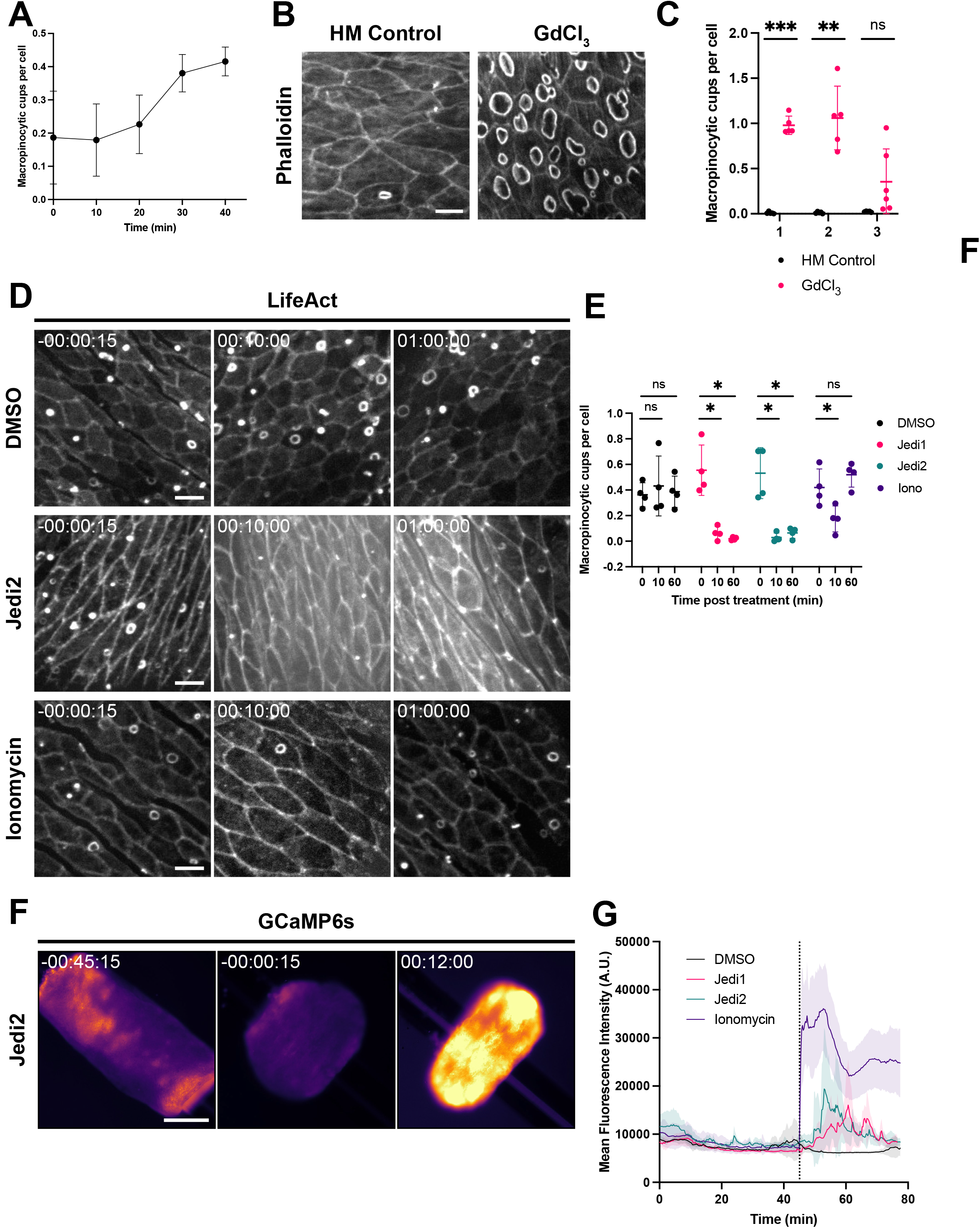
Stretch-activated channel activity regulates macropinocytosis. A) Quantification of macropinocytic cup abundance over time in threaded body columns (mean ± sd; n = 4 independent sample preparations). t = 0 min corresponds to the earliest acquirable time point after body column amputation and threading. B) Representative images of macropinocytic cups stained with phalloidin in *Hydra* medium (HM) control (left) and GdCl_3_-treated (right; 50 μM for 15 min) intact *Hydra.* Scale bar, 20 μm. C) Quantification of macropinocytic cup frequency in response to specified treatments shown in (B). 3 Independent sample preparations are shown per condition (Bars: mean ± sd; n = 5 animals per condition per independent sample preparation; ns = not significant, ** = p < 0.01, *** = p < 0.001, Student’s t-test). D) Representative timecourses of threaded body columns expressing LifeAct-GFP in the ectoderm before (left) and after (center, right) treatment with DMSO, Jedi2 (200 μM), or ionomycin (10 μM). Scale bars, 20 μm. E) Quantification of macropinocytic cup frequency before, 10 min after, and 1 h after specified treatments shown in (D) (Bars: mean ± sd; n = 4 independent sample preparations per condition; ns = not significant, * = p < 0.05, Student’s t-test). See also Figure S2C. F) Representative time-course of a threaded body column expressing GCaMP6s in the ectoderm, before (left, center) and after (right) Jedi2 treatment (200 μM). Images represent single z-sections. mpl-inferno LUT (FIJI) was applied to aid in visualizing graded GCaMP6s signal. Scale bar, 500 μm. G) Traces depicting mean fluorescence intensity over time in threaded body columns before and after specified treatment (line and shading: mean ± sd; n = 4 independent sample preparations per condition). Dotted line indicates drug or vehicle addition. See also Figure S2D. (B, D) All frames depict maximum intensity projections of 15–35 μm z-stacks. (D, F) Time stamps indicate time relative to drug addition; hh:mm:ss.

Stretch-activated channels play a role in tension sensing in a variety of biological contexts [22], and analysis of recent *Hydra* single-cell RNA sequencing data [23] revealed that the *Hydra* ectoderm expresses several putative TRP (e.g. t27236aep, t29007aep) and Piezo (t21136aep) channel proteins that may contribute to mechanosensation. To determine if stretch-activated channels modulate macropinocytic cup formation, we treated *Hydra* with gadolinium chloride (GdCl_3_), a broad-spectrum inhibitor of these channels [24]. Immunofluorescence and live imaging in intact animals revealed a striking increase in macropinocytic cups following treatment with GdCl_3_ (50 μM; Fig. 2B, 2C, S2A). We observed no significant increase in the frequency of macropinocytic cups in GdCl_3_-treated threaded body columns over their already elevated levels of macropinocytosis (Fig. S2B; Video S2B), suggesting that the effect observed in this preparation may indeed result from mechanical unloading. To further probe the role of stretch-activated channels, we performed the reciprocal experiment of treating *Hydra* body columns with Jedi1 and Jedi2, which are activators of the stretch-activated channel Piezo1 [25]. Treatment with either Jedi drug resulted in a near complete, albeit transient, depletion of macropinocytic cups in isolated body columns, coinciding with contraction of the tissue (Fig. 2D, 2E, S2C; Video S3A-C). Together, these findings implicate stretch-activated channels, including Piezo1, in the regulation of macropinocytosis in *Hydra*.

Upon activation, stretch-activated channels signal mechanical stimuli through the transport of ions, predominantly Ca^2+^ influx in the case of Piezo channels [26]. To directly monitor calcium flux in the *Hydra* ectoderm, we prepared threaded body columns from transgenic *Hydra* expressing the fluorescent calcium reporter GCaMP6s in ectodermal tissues [27]. Intriguingly, we observed a gradual decrease in fluorescence intensity as body columns conformed to their filaments (Fig. 2F, 2G, S2D; Video S3E-G), coinciding with the period of increasing macropinocytosis reported above. Treatment of body columns with Jedi1 or Jedi2 induced a transient spike in GCaMP6s fluorescence, accompanied by strong contractions (Fig. 2F, 2G, S2D; Video S3E-G).

To further test the extent to which Jedi1/2 inhibited macropinocytosis via changes in intracellular calcium, we treated threaded *Hydra* body columns with ionomycin to promote stretch-activated channel-independent calcium influx. Using GCaMP6s-expressing *Hydra*, we confirmed that ionomycin induced an increase in fluorescence intensity and body column contractions, indicating an increase in cytosolic calcium concentrations (Fig. 2G, S2D; Video S3H). In LifeAct-GFP-expressing body columns, ionomycin treatment resulted in a transient decrease in macropinocytic cups. Intriguingly, this effect was less pronounced and shorter lived than that observed upon exposure to Jedi1 or Jedi2 (Fig. 2D, 2E; Video S3D), despite a greater and more sustained calcium influx following ionomycin treatment (Fig. 2G). Thus, while our findings suggest that calcium influx is sufficient to transiently inhibit macropinocytosis, this alone may not explain the magnitude of the effect observed following Piezo1 activation.

### Mechanical stretch inhibits macropinocytosis

In light of our data implicating stretch-activated channels in regulating macropinocytosis, we sought to directly test the effects of tissue stretch on this process. Body column fragments that are allowed to heal (i.e. without threading onto filaments) form hollow tissue spheres, which undergo cycles of swelling and rupturing and are capable of ultimately regenerating healthy *Hydra* [28,29]. We sought to capitalize on the architectural simplicity and deformability of these regenerating “spheroids” as a system for applying stretch. To this end, we microinjected *Hydra* medium into the lumen of *Hydra* spheroids, inflating them to approximately 1.5 times their original volume before they ruptured (approximated volume fold change: 1.47 ± 0.19; Fig. 3A, S3; Video S4A). In an inflated state, ectodermal cells of LifeAct-GFP-expressing spheroids exhibited enlarged apical surface areas, suggestive of a planar ectodermal stretch (Fig. 3B).

**Figure 3.**
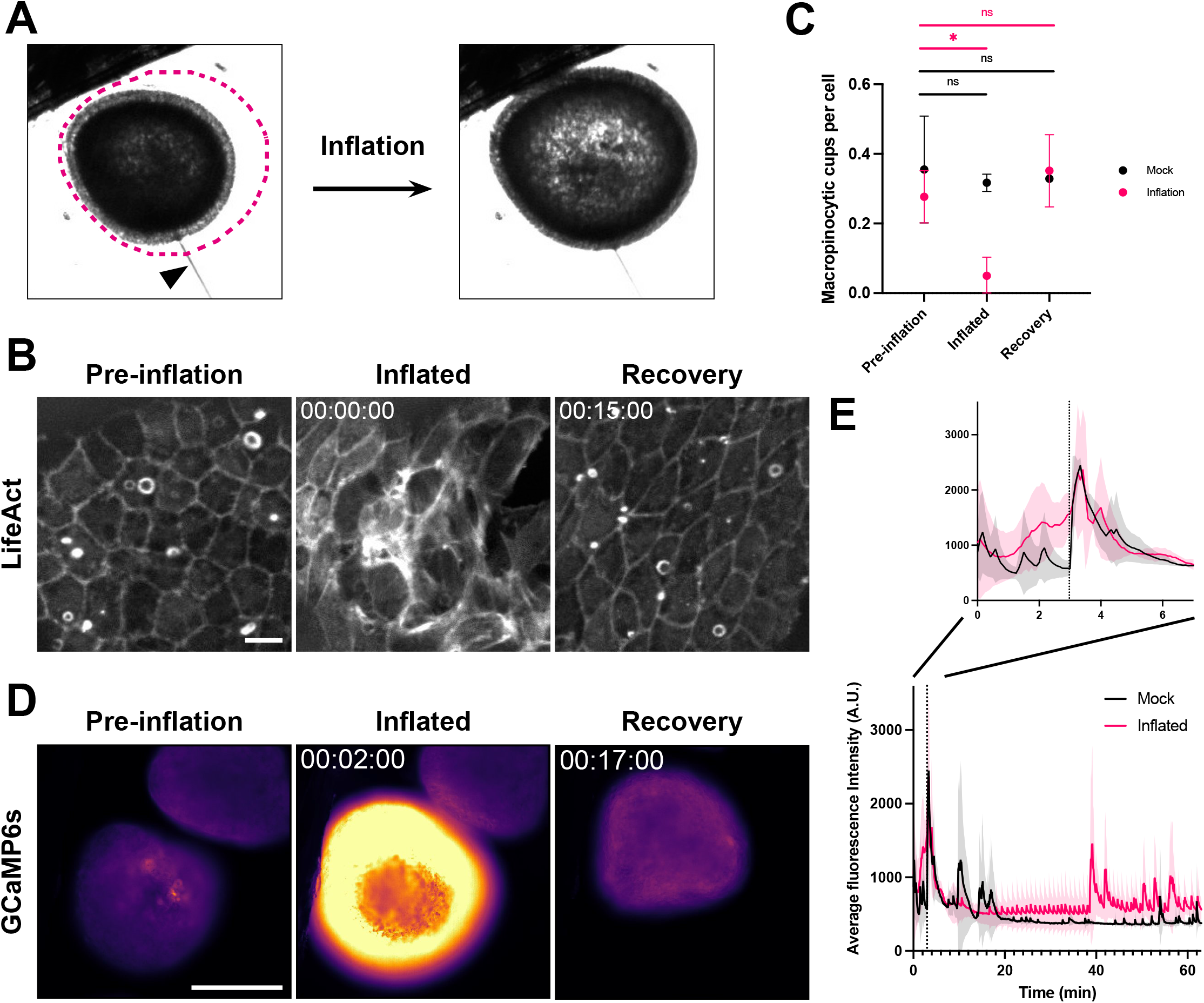
Application of tissue stretch inhibits macropinocytosis. A) Representative images of the same *Hydra* spheroid before (left) and after inflation (right). Magenta trace indicates the profile of the spheroid at peak inflation. Arrowhead: microinjection needle. B) Representative time-course of the same spheroid expressing ectodermal LifeAct-GFP before (Pre-inflation), immediately after inflation and needle removal (Inflated), and during recovery (Recovery). Time stamp indicates time relative to needle removal following inflation; hh:mm:ss. All frames depict maximum intensity projections of 15–35 μm z-stacks. Pre-inflation and Inflated/Recovery images have been scaled independently to compensate for an increase in signal following inflation. Scale bar, 20 μm. C) Quantification of macropinocytic cup frequency during inflation experiments shown in (B) (mean ± sd; n = 3 independent sample preparations per condition; ns = not significant, * = p < 0.05, Student’s t-test). D) Representative time-course of the same spheroid expressing ectodermal GCaMP6s before (Pre-inflation), immediately after inflation (Inflated), and during recovery (Recovery). mpl-inferno LUT (FIJI) was applied to aid in visualizing graded GCaMP6s signal. Time stamp indicates time relative to initiation of inflation; hh:mm:ss. All frames depict single z-sections. Scale bar, 200 μm. E) Traces depicting mean GCaMP6s fluorescence intensity over time during inflation experiments (line and shading: mean ± sd; n = 3 independent sample preparations per condition). Dotted line indicates needle removal.

We next characterized the abundance of macropinocytic cups in inflated and uninflated LifeAct-GFP-expressing spheroids. Macropinocytic cups were depleted in inflated aggregates when compared to their pre-inflation states. As aggregates gradually deflated following needle removal, we observed a recovery of macropinocytosis (Fig. 3B, 3C; Video S4C). Applying the same protocol to GCaMP6s-expressing spheroids, we observed a dramatic increase in GCaMP6s fluorescence during spheroid inflation. As spheroids deflated, GCaMP6s fluorescence gradually diminished to basal levels (Fig. 3D, 3E; Video S4D). Needle insertion and removal also resulted in brief spikes in GCaMP6s intensity (Fig. 3E; Video S4D). However, mock inflations, in which a microinjection needle was inserted into spheroids for a comparable duration without inflation, did not significantly affect macropinocytosis (Fig. 3C; Video S4B). Together these data indicate that inflation of spheroids, with an accompanying stretch of the epithelial layer, leads to calcium influx and inhibits macropinocytosis.

## Discussion

Here, we describe widespread macropinocytosis in the outer epithelium of *Hydra vulgaris*. Our results suggest that this phenomenon is regulated by tension applied to the epithelial layer. This is most directly demonstrated by the finding that inducing stretch in *Hydra* spheroids through inflation is sufficient to transiently inhibit macropinocytosis, with macropinocytosis rapidly recovering following deflation and relaxation of spheroids. This tissue stretch coincides with a rise in intracellular calcium in epithelial cells. We further show that pharmacological inhibition of ion flux through stretch-activated channels results in the activation of macropinocytosis and that Piezo1 activation or calcium ionophores repress macropinocytosis. Together, our findings highlight a role for tissue mechanics and stretch-activated channels in the regulation of macropinocytosis in *Hydra* (Fig. 4).

**Figure 4.**
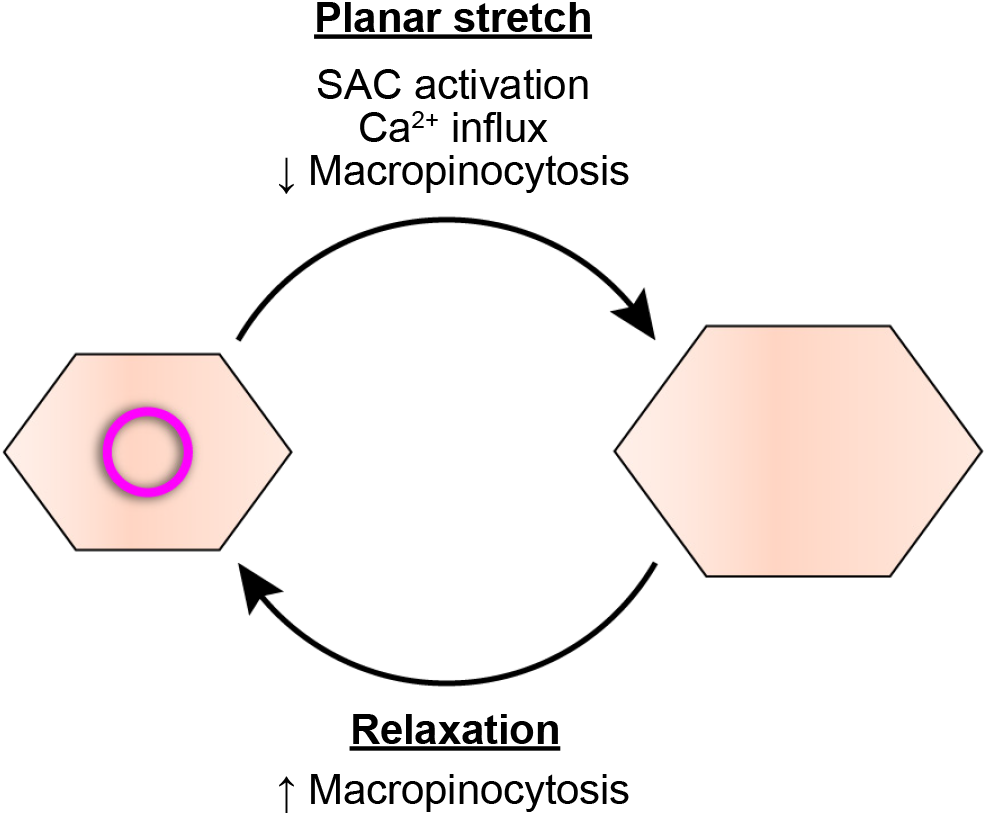
Mechanical stretch-regulated macropinocytosis. Model of mechanically regulated macropinocytosis in *Hydra*. Planar stretch results in stretch-activated channel (SAC) activation, calcium influx, and inhibition of macropinocytosis. Following tissue relaxation, SAC inactivation restores macropinocytosis.

Macropinocytosis is historically categorized as “constitutive” or “stimulated,” based on the requirement for growth factors for ruffle induction. Constitutive macropinocytosis has been attributed to relatively few cell types, most notably, amoebae and antigen presenting cells, where respective roles in feeding and immune surveillance have been described (reviewed in [30,31]). In contrast, most mammalian epithelial models for which macropinocytosis has been reported require growth factor stimulation. Given that macropinocytosis in *Hydra* occurs in a wide variety of culture conditions without supplementation, our findings likely reflect an underappreciated form of constitutive epithelial macropinocytosis. Intriguingly, a similar ubiquitous macropinocytosis was recently reported in the tissues of other cnidarian species [32], raising the possibility that constitutive macropinocytosis is more widespread than previously appreciated, at least among basal metazoans.

Our findings directly implicate membrane tension as a regulator of macropinocytosis, expanding the repertoire of exogenous effectors beyond growth factors [12] and pathogens [11]. Precisely how properties like membrane tension may contribute to macropinocytosis remains unresolved. Membrane tension may be transduced to biochemical signals that limit actin nucleation and membrane protrusion. A similar mechanical-biochemical crosstalk has been described in the actin-mediated migration of neutrophils [33,34]. Alternatively, macropinocytic cups may form and evolve more readily in low-tension conditions, as the energetic cost to deform membranes may be reduced [35]. While we cannot exclude the latter possibility, our finding that the pharmacological inhibition of stretch-activated channels is sufficient to induce macropinocytosis suggests an active contribution.

In our experiments, gadolinium, which broadly inhibits stretch-activated channels, promoted macropinocytosis. Moreover, application of the Piezo-specific activators Jedi1/2 [25] resulted in a transient inhibition of macropinocytosis, accompanied by calcium influx and tissue contraction. This indicates a role for, but not limited to, Piezo channels in macropinocytosis regulation. Piezo function has been extensively explored in epithelia, where Piezo1 indirectly senses cellular crowding via membrane tension to maintain homeostatic cell densities [36,37]. Intriguingly, recent work similarly identified Piezo1 as a regulator of epidermal growth factor-stimulated macropinocytosis in cancer cells [38]. In mammalian myotubes, where a role for membrane tension in macropinocytosis has also been reported, tension-mediated redistribution of phospholipase D2 in the plasma membrane was proposed to promote actin remodeling and membrane ruffling, suggesting an alternative mechanisms to transduce mechanical stimuli to biochemical signals [14,15]. Thus, tissue stretch may play a role in macropinocytosis in a number of contexts, and whether additional factors beyond stretch-activated channels contribute to this process in *Hydra* remains to be determined.

Stretch-activated channels provide a means to transduce mechanical stimuli to chemical signals by conducting ions across the plasma membrane [26]. Using ionomycin to induce calcium influx independent of stretch-activated channels, we observed a lesser and more transient inhibition of macropinocytosis when compared to treatment with Piezo1 activators Jedi1/2. Piezos are non-selective cation channels permeable to a number of ions in addition to Ca^2+^, including K^+^, Na^+^, and Mg^2+^, all of which are present in *Hydra* medium and may, therefore, explain some of the observed effect of Piezo activation [39]. Alternatively, it is conceivable that the robust and sustained depolarization that occurs in the presence of ionomycin interferes with the mechanisms required to inhibit macropinocytosis. Future studies will provide further insight into the requirements for macropinocytosis and its inhibition in *Hydra*.

Although both our pharmacological and physical perturbations indicate a role for stretch in inhibiting macropinocytosis, we note several reasons for caution in interpreting these findings. For instance, calcium influx causes contraction of *Hydra*’s epithelial myonemes, which may, in turn, alter tension at the apical membrane. Thus, while Jedi1/2 and ionomycin treatment transiently inhibited macropinocytosis, these effects may be an indirect consequence of contraction, rather than a direct effect of Piezo activation. Further genetic analysis will be useful in dissecting the mechanistic roles of Piezo channels and calcium signaling in macropinocytosis in *Hydra*. Additionally, amputated and isolated body columns provide an unparalleled opportunity to observe macropinocytosis but require substantial tissue wounding, which may contribute to the influx of macropinocytic cups we observe in this context. Nevertheless, our spheroid inflation approach circumvents acute injury on this scale and provides the most direct evidence for tissue mechanics in the regulation of macropinocytosis.

The role of macropinocytosis in *Hydra* remains unknown. Given *Hydra*’s predatory feeding behaviors, macropinocytosis is unlikely to play a significant role in nutrient acquisition. Similarly, the dilute solutes available in *Hydra*’s freshwater habitats likely render this process inefficient for acquiring or responding to dissolved environmental factors. Alternatively, macropinocytosis may contribute to the uptake and remodeling of contents already associated with the animal’s surface. Notably, *Hydra* epithelia have been shown to internalize particles of various sizes, including beads of up to 1 μm in diameter [40,41], which demonstrates an ability to indiscriminately engulf solid substrates that may be explained by our findings. In one study [40], ultrastructural analysis revealed large, nanoparticle-filled vacuoles containing remains of the *Hydra* “cuticle,” a fibrous structure secreted by and surrounding *Hydra*’s apical ectodermal surface and home to a complex microbial community [42–44]. Thus, macropinocytosis may provide an efficient means to remodel the *Hydra* cuticle and sense and respond to the microbiome.

Given our data showing stretch-sensitivity, we also speculate that macropinocytosis could play a role in regulating membrane tension through the removal of large regions of the apical plasma membrane. The *Hydra* epithelium is a remarkably dynamic tissue, characterized by perpetual growth and cell loss, extensive cellular rearrangements ([45]; reviewed in [46]), and drastic changes in aspect ratio during animal contraction and elongation. Based on measurements of macropinocytic cup size and macropinosome accumulation, we estimate rates of membrane retrieval in isolated body columns exceeding 150 μm^2^ per min per 10,000 μm^2^, corresponding to complete apical membrane turnover in approximately 1 h. Thus, our findings suggest a significant capacity for membrane recycling through macropinocytosis. Together with the regulatory role for cellular stretch that we identify, it is tempting to speculate that this capacity to rapidly recycle apical membrane may contribute to maintaining membrane tension and remodeling tissues. Although we did not observe direct evidence for such a mechanism, this may be confounded by comparable rates of membrane deposition in isolated body columns, as we observed no apparent decreases in apical membrane surface area during this process. Future efforts to uncouple membrane retrieval and deposition may shed light on a role for macropinocytosis in membrane tensioning and, more broadly, reveal the contributions of tension-regulated macropinocytosis to *Hydra* physiology.

## Acknowledgments

We thank Celina Juliano and Rob Steele for numerous insights into *Hydra* care and experimentation. We are grateful to Rob Steele and Rafael Yuste for providing additional *Hydra* lines and reagents. This material is based upon work supported by the National Science Foundation Graduate Research Fellowship under Grant No. 1650113 (to TDS). TDS was also supported by the UCSF Chuan Lyu Discovery Fellowship. BH was supported by the Marie Skłodowska-Curie COFUND program ARDRE “Ageing, Regeneration and Drug Research” (Grant No. 847681). KLM was supported by the Damon Runyon Cancer Research Foundation (DRG-2282-17 and DFS-47-21) and by the Eunice Kennedy Shriver National Institute Of Child Health & Human Development of the National Institutes of Health (K99HD101021). RDV was supported by the Howard Hughes Medical Institute.

## Author Contributions

Investigation, Methodology, Validation, Formal Analysis, Visualization: T.D.S.; Writing-original draft: T.D.S., K.L.M; Supervision: K.L.M., R.D.V.; Conceptualization, Writing–review and editing, Funding Acquisition: T.D.S., K.L.M., R.D.V., B.H..

## Declaration of Interests

The authors declare no competing interests.

## Materials and methods

### Hydra culturing and strains

*Hydra* were maintained at 18 °C in *Hydra* medium and fed 2-3 times per week with *Artemia* nauplii (Brine Shrimp Direct). Animals were starved ≥ 24h prior to experimentation, and nonbudding animals were chosen for experimentation. The following transgenic lines were used:

DsRed2(ectoderm)/GFP(endoderm) [47]
LifeAct-GFP(ectoderm) [17]
pActin::GCaMP6s(ectoderm) [27]
AEP SS1 (courtesy of Rob Steele)

### Microscopy

Immunofluorescence and live images of LifeAct-expressing spheroids and body columns were acquired on a Yokogawa CSU100 spinning disk confocal attached to an inverted Nikon Ti-E microscope, with Hamamatsu C9100-13 EMCCD camera, using 20X Plan Apo VC 0.75 NA or 60XA Plan Apo VC 1.20 NA WI objectives. Images of GCaMP6s-expressing spheroids were acquired on an inverted Nikon Ti-E microscope with Lumencor SpectraX epifluorescence module and Andor Zyla camera, using a 10X Plan Apo 0.45 NA objective. Images of GCaMP6s-expressing body columns were acquired on an inverted Zeiss Axiovert 200M microscope with and Point Grey Chamelion3 Monochrome camera, using a 5X EC Plan-Neofluar 0.16 NA objective. Confocal Z-stacks were acquired at 2-10 μm step sizes for a total depth of 30-100 μm, at 4-60 s time intervals. Where epifluorescence was used, images were acquired at a fixed focal plane at 5-30 s intervals. All images were acquired using Micro-Manager software [48].

### Immunofluorescence

Immunofluorescence was performed as previously described [17], with slight modification. In brief, AEP SS1 *Hydra* were relaxed for 1 min in 2% urethane (Sigma, Cat# 51-79-6)/*Hydra* medium, fixed for 1 h in 4% PFA (Electron Microscopy Sciences, Cat# 15714)/*Hydra* medium, washed 3x in PBS (Gibco, Cat# 20012-027), permeabilized for 15 min in 0.5% Triton X-100 (Sigma, Cat# 9036-19-5)/PBS, blocked for 1 h in 1% BSA (Sigma, Cat# A7906)/0.1% Triton X-100/PBS (blocking solution), stained for 1 h in Alexa 488-phalloidin (Invitrogen, Cat# A12379) diluted to 1:200 in blocking solution, washed 3x in PBS, and mounted between a glass slide and coverslip with ProLong Gold mounting medium (Invitrogen, Cat# P36930).

### Tissue manipulations

Threaded body columns were prepared by amputating the head and foot (peduncle) from *Hydra* with a scalpel and inserting a 5-10 mm length of 8-lb fishing line (Trilene SensiThin) through the exposed lumen. Body columns were allowed to stabilize for 15 min prior to chemical perturbations, but were otherwise imaged immediately after preparation and transfer to imaging vessels, which consisted of either 35mm glass bottom dishes (MatTek, P35G-1.5-14-C) or 96-well glass bottom plates (MatriPlate, MGB096-1-2-LG-L).

Spheroids were prepared by removing the head and foot from *Hydra* and cutting the remaining body column into 3-4 rings, which were subsequently cut longitudinally into 2-3 sections. Dissected tissues were allowed to heal unperturbed for 12-24 h prior to experimentation. Micro-injection/inflation was achieved using a micropipette mounted to a Narishige motor-driven micro-manipulator (MM-94) via Narishige microscope mounting adaptor, injection holder, and universal joint (NN-H-4, HI-9, UT-2, respectively). Pipettes were pulled from Sutter Instrument capillary tubes (#B150-110-10) on a Sutter Instrument micropipette puller (P-1000). Fluid ejection was controlled by a syringe attached to the pipette. Inflations were performed over a period of 2-3 minutes, after which the micropipette was immediately removed. Only spheroids that remained intact (did not rupture) were used to quantify macropinocytic cup abundance/dynamics. For mock inflations, a micropipette was inserted into the spheroid for 3 minutes, without injection, before removal.

### Dextran uptake

Macropinosomes and dextran uptake were visualized by transferring stabilized threaded body columns to a solution containing pHrodo Red Dextran, 10,000 MW (ThermoFisher, Cat# P10361) diluted to 5 μg/mL in *Hydra* medium, and immediately beginning acquisitions.

### Chemical perturbations

GdCl_3_ (Sigma, Cat# 439770) stock solution was prepared in *Hydra* medium and diluted to a final concentration of 50 μM. For immunofluorescence experiments, animals were directly transferred to either the 50 μM GdCl_3_ solution or fresh *Hydra* medium (HM control) and incubated for 15 min prior to fixation. For threaded body column experiments, GdCl_3_ was diluted directly into imaging chamber *Hydra* medium to a final concentration of 50 μM. For drug perturbations, Jedi1 (Sigma, SML2533), Jedi2 (Sigma, SML2532), and ionomycin calcium salt (Sigma, I0634) stock solutions were prepared in DMSO and diluted directly into imaging chamber *Hydra* medium to final concentrations of 200 μM, 200 μM, and 10 μM, respectively. DMSO controls corresponded to the highest DMSO concentration (1% v/v) present in drug treatments.

### Quantification and Statistical Analysis

Measurements of macropinocytic cup sizes were performed in FIJI using the built-in measure function for a line segment drawn along the long axis of macropinocytic cups at their maximum width. Measures of macropinocytic cups per cell were obtained from manual counts of macropinocytic cups and cells occupying the microscope field of view at a given time point. Quantifications assigned to the head, body column, and foot were generated from images obtained from the top, middle, and bottom 1/3 of the animal’s body length (tentacles excluded). Quantification of macropinosomes was similarly obtained by manual counts of dextran-filled puncta at a given time point. Average GCaMP6s fluorescence intensity was quantified by generating binary masks corresponding to body columns or spheroids in each frame, applying masks to unadulterated images to define regions of interest (ROIs), and quantifying the mean gray value of all pixels within the specified ROI using FIJI’s built-in measure function. The estimated fold change in spheroid volume before rupture was determined by approximating each spheroid as a true sphere and using the manually measured radius along the spheroid long axis before and after inflation (until rupture) for calculations.

## Supplemental Figure Legends

**Figure S1.** Schematic (left), representative images (middle), and quantification (right) of macropinocytic cup abundance in the head, body column, and foot of fixed, intact *Hydra* stained with phalloidin (mean ± sd; n ≥ 18 animals from 3 independent sample preparations per body region). All frames depict maximum intensity projections of 10-35 μm z-stacks.

**Figure S2.** A) Representative images of live, intact *Hydra* expressing LifeAct-GFP in the ectoderm, treated with control *Hydra* medium (left) or GdCl_3_ (right; 50 μM for 15 min). B) Quantification of macropinocytic cup frequency in *Hydra* medium control (black) and GdCl_3_-treated (magenta; 50 μM) threaded body columns over time (mean ± sd; n = 4 independent sample preparations per condition). Note that HM control data is duplicated from Fig. 2A. C) Representative time-course of a threaded body column expressing LifeAct-GFP in the ectoderm before (left) and after (center, right) treatment with Jedi1 (200 μM). (A, C) Scale bars, 20 μm. All frames depict maximum intensity projections of 15-35 μm z-stacks. D) Representative time-courses of threaded body columns expressing GCaMP6s in the ectoderm, before (left, center) and after (right) DMSO, Jedi1 (200 μM), or Ionomycin (10 μM) treatment. Images depict single z-sections. Scale bars, 500 μm. (C, D) Time stamps indicate time relative to drug addition; hh:mm:ss.

**Figure S3.** Quantification of the approximated fold change in spheroid volume achievable prior to tissue rupture during spheroid inflation (bars: mean ± sd; n = 11 from 4 independent sample preparations).

## Supplemental Video Legends

**Video S1.** A) Representative time-course of macropinocytic cup formation, closure, and dissipation, visualized by LifeAct-GFP. Image registration was performed to compensate for translational movement in the body column. Corresponds to Figure 1D. B) Representative timecourse of fluorescently labeled dextran (magenta) engulfment by macropinocytic cups, visualized by LifeAct-GFP (white). Image registration was performed to compensate for translational movement in the body column. Corresponds to Figure 1E. (A, B) All videos depict maximum intensity projections of 10–50 μm z-stacks. Time stamps, hh:mm:ss.

**Video S2.** A) Representative time-course of macropinocytic cup frequency in a control isolated body column treated with *Hydra* medium, visualized by LifeAct-GFP. Corresponds to Figure 2A. B) Representative time-course of macropinocytic cup frequency in a GdCl_3_-treated (50 μM) isolated body column, visualized by LifeAct-GFP. Corresponds to Figure S2B. (A, B) All videos depict maximum intensity projections of 35–100 μm z-stacks. Time stamps indicate time relative to drug addition; hh:mm:ss.

**Video S3.** A) Representative time-course of macropinocytic cup frequency in a control isolated body column treated with DMSO, visualized by LifeAct-GFP. B) Representative time-course of macropinocytic cup frequency in a Jedi2-treated (200 μM) isolated body column, visualized by LifeAct-GFP. C) Representative time-course of macropinocytic cup frequency in a Jedi1-treated (200 μM) isolated body column, visualized by LifeAct-GFP. D) Representative time-course of macropinocytic cup frequency in an ionomycin-treated (10 μM) isolated body column, visualized by LifeAct-GFP. (A-D) Correspond to Figures 2D, 2E, S2C. All videos depict maximum intensity projections of 35-100 μm z-stacks. E) Representative time-course of calcium signaling in a control isolated body column treated with DMSO, visualized by GCaMP6s. F) Representative time-course of calcium signaling in a Jedi2-treated (200 μM) isolated body column, visualized by GCaMP6s. G) Representative time-course of calcium signaling in a Jedi1-treated (200 μM) isolated body column, visualized by GCaMP6s. H) Representative time-course of calcium signaling in an ionomycin-treated (10 μM) isolated body column, visualized by GCaMP6s. (E-H) Correspond to Figures 2F, 2G, S2D. All videos depict single z-sections. (A-H) Time stamps indicate time relative to drug addition; hh:mm:ss.

**Video S4.** A) Representative time-course of spheroid inflation. Corresponds to Figure 3A. Video depicts a single z-section. B) Representative time-course of macropinocytic cup frequency in the same control spheroid imaged before (pre-mock inflation) and after (post-mock inflation) mock inflation, visualized by LifeAct-GFP. Note “pre-mock inflation” depicts a still image (single time point). C) Representative time-course of macropinocytic cup frequency in the same inflated spheroid imaged before (pre-inflation) and after (post-inflation) inflation, visualized by LifeAct-GFP. Note “pre-inflation” depicts a still image (single time point). (B, C) Correspond to Figures 3B, 3C. All videos depict maximum intensity projections of 35-100 μm z-stacks. Time stamp indicates time relative to needle removal following treatment; hh:mm:ss. D) Representative timecourse of calcium signaling in spheroid during and after inflation and needle removal, visualized by GCaMP6s. “INFLATE” label marks period of inflation. Note the second calcium transient upon needle removal, immediately after inflation period. Corresponds to Figures 3D, 3E. Video depicts a single z-section. Time stamp indicates time relative to initiation of inflation; hh:mm:ss.

## Notes

### Competing Interest Statement

The authors have declared no competing interest.

## References

1. Kerr, M.C., and Teasdale, R.D. (2009). Defining macropinocytosis. Traffic 10, 364–371.

2. Bloomfield, G., and Kay, R.R. (2016). Uses and abuses of macropinocytosis. Journal of Cell Science 129, 2697–2705.

3. Bloomfield, G., Traynor, D., Sander, S.P., Veltman, D.M., Pachebat, J.A., and Kay, R.R. (2015). Neurofibromin controls macropinocytosis and phagocytosis in Dictyostelium. eLife 4, e04940.

4. Commisso, C., Davidson, S.M., Soydaner-Azeloglu, R.G., Parker, S.J., Kamphorst, J.J., Hackett, S., Grabocka, E., Nofal, M., Drebin, J.A., Thompson, C.B., et al. (2013). Macropinocytosis of protein is an amino acid supply route in Ras-transformed cells. Nature 497, 633–637.

5. Sallusto, F., Cella, M., Danieli, C., and Lanzavecchia, A. (1995). Dendritic cells use macropinocytosis and the mannose receptor to concentrate macromolecules in the major histocompatibility complex class II compartment: Downregulation by cytokines and bacterial products. Journal of Experimental Medicine 182, 389–400.

6. Sarkar, K., Kruhlak, M.J., Erlandsen, S.L., and Shaw, S. (2005). Selective inhibition by rottlerin of macropinocytosis in monocyte-derived dendritic cells. Immunology 116, 513–524.

7. Orth, J.D., Krueger, E.W., Weller, S.G., and McNiven, M.A. (2006). A novel endocytic mechanism of epidermal growth factor receptor sequestration and internalization. Cancer Res 66, 3603–3610.

8. Clayton, E.L., Evans, G.J.O., and Cousin, M.A. (2008). Bulk synaptic vesicle endocytosis is rapidly triggered during strong stimulation. J. Neurosci. 28, 6627–6632.

9. Holt, M., Cooke, A., Wu, M.M., and Lagnado, L. (2003). Bulk membrane retrieval in the synaptic terminal of retinal bipolar cells. J Neurosci 23, 1329–1339.

10. Buckley, C.M., and King, J.S. (2017). Drinking problems: Mechanisms of macropinosome formation and maturation. The FEBS Journal 284, 3778–3790.

11. Rosales-Reyes, R., Pérez-López, A., Sánchez-Gómez, C., Hernández-Mote, R.R., Castro-Eguiluz, D., Ortiz-Navarrete, V., and Alpuche-Aranda, C.M. (2012). Salmonella infects B cells by macropinocytosis and formation of spacious phagosomes but does not induce pyroptosis in favor of its survival. Microbial Pathogenesis 52, 367–374.

12. West, M.A., Bretscher, M.S., and Watts, C. (1989). Distinct endocytotic pathways in epidermal growth factor-stimulated human carcinoma A431 cells. Journal of Cell Biology 109, 2731–2739.

13. Chiasson-MacKenzie, C., Morris, Z.S., Liu, C.-H., Bradford, W.B., Koorman, T., and McClatchey, A.I. (2018). Merlin/ERM proteins regulate growth factor-induced macropinocytosis and receptor recycling by organizing the plasma membrane:cytoskeleton interface. Genes Dev. Available at: http://genesdev.cshlp.org/content/early/2018/08/24/gad.317354.118 [Accessed September 3, 2018].

14. Lin, S.-S., and Liu, Y.-W. (2019). Mechanical stretch induces mTOR recruitment and activation at the phosphatidic acid-enriched macropinosome in muscle cell. Front Cell Dev Biol 7. Available at: https://www.ncbi.nlm.nih.gov/pmc/articles/PMC6518962/ [Accessed June 10, 2021].

15. Loh, J., Chuang, M.-C., Lin, S.-S., Joseph, J., Su, Y.-A., Hsieh, T.-L., Chang, Y.-C., Liu, A.P., and Liu, Y.-W. (2019). An acute decrease in plasma membrane tension induces macropinocytosis via PLD2 activation. Journal of Cell Science 132. Available at: https://doi.org/10.1242/jcs.232579 [Accessed May 28, 2021].

16. Lin, X.P., Mintern, J.D., and Gleeson, P.A. (2020). Macropinocytosis in different cell types: Similarities and differences. Membranes 10, 177.

17. Aufschnaiter, R., Wedlich-Söldner, R., Zhang, X., and Hobmayer, B. (2017). Apical and basal epitheliomuscular F-actin dynamics during Hydra bud evagination. Biol Open 6, 1137–1148.

18. Skokan, T.D., Vale, R.D., and McKinley, K.L. (2020). Cell sorting in Hydra vulgaris arises from differing capacities for epithelialization between cell types. Current Biology 30, 3713–3723.e3.

19. Beams, H.W., Kessel, R.G., and Shih, C.-Y. (1973). The surface features of Hydra as revealed by scanning electron microscopy. Transactions of the American Microscopical Society 92, 161.

20. Mellström, K., Höglund, A.-S., Nistér, M., Heldin, C.-H., Westermark, B., and Lindberg, U. (1983). The effect of platelet-derived growth factor on morphology and motility of human glial cells. J Muscle Res Cell Motil 4, 589–609.

21. Buccione, R., Orth, J.D., and McNiven, M.A. (2004). Foot and mouth: Podosomes, invadopodia and circular dorsal ruffles. Nature Reviews Molecular Cell Biology 5, 647–657.

22. Stewart, T.A., and Davis, F.M. (2019). Formation and function of mammalian epithelia: Roles for mechanosensitive PIEZO1 ion channels. Frontiers in Cell and Developmental Biology 7, 260.

23. Siebert, S., Farrell, J.A., Cazet, J.F., Abeykoon, Y., Primack, A.S., Schnitzler, C.E., and Juliano, C.E. (2019). Stem cell differentiation trajectories in Hydra resolved at single-cell resolution. Science 365, eaav9314.

24. Yang, X., and Sachs, F. (1989). Block of stretch-activated ion channels in Xenopus oocytes by gadolinium and calcium ions. Science 243, 1068–1071.

25. Wang, Y., Chi, S., Guo, H., Li, G., Wang, L., Zhao, Q., Rao, Y., Zu, L., He, W., and Xiao, B. (2018). A lever-like transduction pathway for long-distance chemical- and mechano-gating of the mechanosensitive Piezo1 channel. Nat Commun 9, 1300.

26. Coste, B., Mathur, J., Schmidt, M., Earley, T.J., Ranade, S., Petrus, M.J., Dubin, A.E., and Patapoutian, A. (2010). Piezo1 and Piezo2 are essential components of distinct mechanically activated cation channels. Science 330, 55–60.

27. Szymanski, J.R., and Yuste, R. (2019). Mapping the whole-body muscle activity of Hydra vulgaris. Current Biology 29, 1807–1817.e3.

28. Sato-Maeda, M., and Tashiro, H. (1999). Development of oriented motion in regenerating Hydra cell aggregates. jzoo 16, 327–334.

29. Shimizu, H., Sawada, Y., and Sugiyama, T. (1993). Minimum tissue size required for Hydra regeneration. Developmental Biology 155, 287–296.

30. King, J.S., and Kay, R.R. (2019). The origins and evolution of macropinocytosis. Philosophical Transactions of the Royal Society B: Biological Sciences 374, 20180158.

31. Liu, Z., and Roche, P.A. (2015). Macropinocytosis in phagocytes: Regulation of MHC class-II-restricted antigen presentation in dendritic cells. Frontiers in Physiology 6, 1.

32. Ganot, P., Tambutté, E., Caminiti-Segonds, N., Toullec, G., Allemand, D., and Tambutté, S. (2020). Ubiquitous macropinocytosis in anthozoans. eLife 9, e50022.

33. Diz-Muñoz, A., Thurley, K., Chintamen, S., Altschuler, S.J., Wu, L.F., Fletcher, D.A., and Weiner, O.D. (2016). Membrane tension acts through PLD2 and mTORC2 to limit actin network assembly during neutrophil migration. PLOS Biology 14, e1002474.

34. Houk, A.R., Jilkine, A., Mejean, C.O., Boltyanskiy, R., Dufresne, E.R., Angenent, S.B., Altschuler, S.J., Wu, L.F., and Weiner, O.D. (2012). Membrane tension maintains cell polarity by confining signals to the leading edge during neutrophil migration. Cell 148, 175–188.

35. Aghamohammadazadeh, S., and Ayscough, K.R. (2009). Under pressure: The differential requirements for actin during yeast and mammalian endocytosis. Nat Cell Biol 11, 1039–1042.

36. Eisenhoffer, G.T., Loftus, P.D., Yoshigi, M., Otsuna, H., Chien, C.-B., Morcos, P.A., and Rosenblatt, J. (2012). Crowding induces live cell extrusion to maintain homeostatic cell numbers in epithelia. Nature 484, 546–549.

37. Gudipaty, S.A., Lindblom, J., Loftus, P.D., Redd, M.J., Edes, K., Davey, C.F., Krishnegowda, V., and Rosenblatt, J. (2017). Mechanical stretch triggers rapid epithelial cell division through Piezo1. Nature 543, 118–121.

38. Kuriyama, M., Hirose, H., Masuda, T., Shudou, M., Arafiles, J.V.V., Imanishi, M., Maekawa, M., Hara, Y., and Futaki, S. (2021). Piezo1 activation using Yoda1 inhibits macropinocytosis and proliferation of cancer cells. bioRxiv, 2021.05.14.444123.

39. Gnanasambandam, R., Bae, C., Gottlieb, P.A., and Sachs, F. (2015). Ionic selectivity and permeation properties of human PIEZO1 channels. PLOS ONE 10, e0125503.

40. Marchesano, V., Hernandez, Y., Salvenmoser, W., Ambrosone, A., Tino, A., Hobmayer, B., M de la Fuente, J., and Tortiglione, C. (2013). Imaging inward and outward trafficking of gold nanoparticles in whole animals. ACS Nano 7, 2431–2442.

41. Technau, U., and Holstein, T.W. (1992). Cell sorting during the regeneration of Hydra from reaggregated cells. Developmental Biology 151, 117–127.

42. Böttger, A., Doxey, A.C., Hess, M.W., Pfaller, K., Salvenmoser, W., Deutzmann, R., Geissner, A., Pauly, B., Altstätter, J., Münder, S., et al. (2012). Horizontal gene transfer contributed to the evolution of extracellular surface structures: The freshwater polyp Hydra is covered by a complex fibrous cuticle containing glycosaminoglycans and proteins of the PPOD and SWT (Sweet Tooth) families. PLOS ONE 7, e52278.

43. Fraune, S., Anton-Erxleben, F., Augustin, R., Franzenburg, S., Knop, M., Schröder, K., Willoweit-Ohl, D., and Bosch, T.C. (2015). Bacteria-bacteria interactions within the microbiota of the ancestral metazoan Hydra contribute to fungal resistance. ISME J 9, 1543–1556.

44. Lentz, T.L. (1964). The cell biology of Hydra. Yale Medicine Thesis Digital Library, 877.

45. Philipp, I., Aufschnaiter, R., Ozbek, S., Pontasch, S., Jenewein, M., Watanabe, H., Rentzsch, F., Holstein, T.W., and Hobmayer, B. (2009). Wnt/β-Catenin and noncanonical Wnt signaling interact in tissue evagination in the simple eumetazoan Hydra. Proceedings of the National Academy of Sciences 106, 4290–4295.

46. Campbell, R.D. (1974). Cell movements in Hydra. American Zoologist 14, 523–535.

47. Glauber, K.M., Dana, C.E., Park, S.S., Colby, D.A., Noro, Y., Fujisawa, T., Chamberlin, A.R., and Steele, R.E. (2013). A small molecule screen identifies a novel compound that induces a homeotic transformation in Hydra. Development 140, 4788–4796.

48. Edelstein, A., Amodaj, N., Hoover, K., Vale, R., and Stuurman, N. (2010). Computer control of microscopes using μManager. Current Protocols in Molecular Biology 92, 14.20.1–14.20.17.

