## Supplementary figures and images for "Mechanical stretch regulates macropinocytosis in *Hydra vulgaris*"

### Supplemental Figures

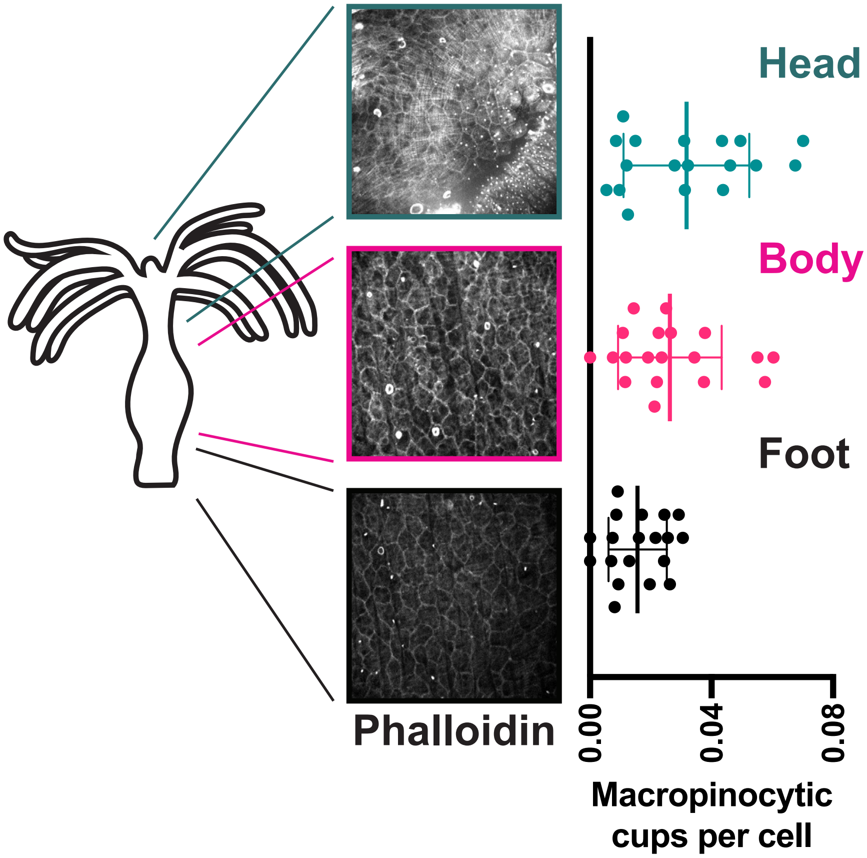

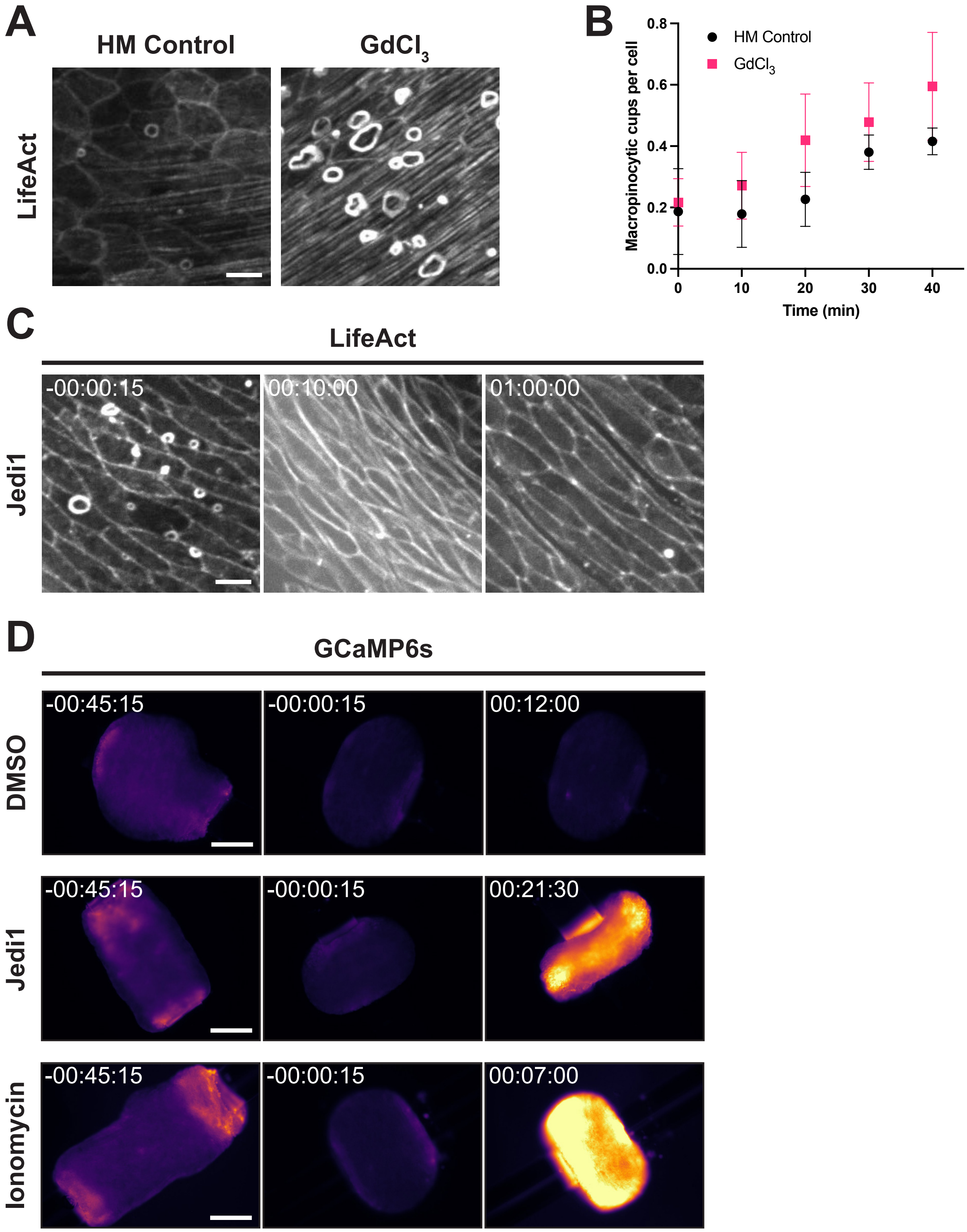

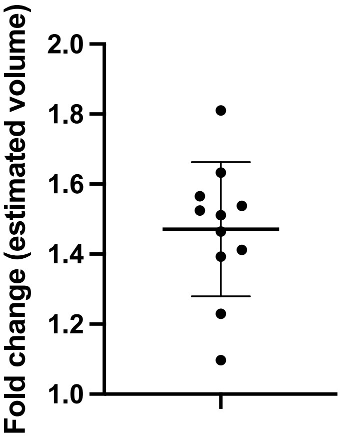


**Figure S3.**
